# Population genomics reveals extensive inbreeding and purging of mutational load in wild Amur tigers

**DOI:** 10.1101/2023.05.09.539923

**Authors:** Tianming Lan, Haimeng Li, Le Zhang, Minhui Shi, Boyang Liu, Liangyu Cui, Nicolas Dussex, Qing Wang, Yue Ma, Dan Liu, Weiyao Kong, Jiangang Wang, Haorong Lu, Shaofang Zhang, Jieyao Yu, Xinyu Wang, Yuxin Wu, Xiaotong Niu, Jiale Fan, Yue Zhao, Love Dalén, Guangshun Jiang, Huan Liu, Yanchun Xu

## Abstract

The inbreeding is a big threat for the persistence of genetic diversity in small and isolated populations of endangered species. The homozygous genome could exacerbate inbreeding depression by introducing homozygous deleterious alleles in the population. However, purging of inbreeding loads as they become homozygotes in small populations could alleviate the depression. The Amur tiger (*Panthera tigris altaica*) is typically exists in small population living in forests in Northeast Asia and is among the most endangered animals on the planet with great symbolic significance of conservation. By comparing with captive individuals, we revealed substantially higher and more extensive inbreeding in the wild Amur tiger population (F_ROH_=0.51) than in captive Amur tigers (F_ROH_=0.26). We further found much less mutational loads in wild populations when compared with captive Amur tigers. However, the frequency of loss of function and deleterious nonsynonymous mutations inside ROH regions are much lower than that in non-ROH regions in both wild and captive Amur tigers, indicating the purging may had occurred in both populations but much effective in the wild population. In addition, we found the average frequency of deleterious alleles was much lower than that of neutral alleles in the wild population, indicating that the purifying selection contributed to the purging of mutational loads in the wild Amur tigers. These findings provide valuable genome-wide evidence to support the making of future conservation plans of wild Amur tigers.

## 1 Introduction

Human-induced habitat fragmentation and environmental changes make a large proportion of biodiversity persisted in small and isolated populations[1-4]. The genetic diversity of a species is of importance for sustaining its evolutionary potentials, which could help it resist the impact from environment changes, thereby reducing the risk of extinction[5, 6]. The genetic viability in small population is highly sensitive to the genetic drift and inbreeding, which could increase the genome homozygosity and then exacerbate the population depression by introducing homozygous mutational loads[7-10]. Genetic rescue is a common strategy to increase genetic diversity and decrease the deleterious effects by building wildlife corridors or reintroduction of captive individuals to the wild to facilitate gene flow[11-13]. However, planning genetic rescue highly depends on the comprehensive investigation of genetic backgrounds, including the inbreeding, outbreeding and introduction of deleterious mutations [14-17], and otherwise may introduce additional genetic loads and/or result in outbreeding depression that further impair the already weak genetic basis.

Purifying selection acts on the small population could help purge deleterious alleles as they become homozygotes, and thereby reducing the inbreeding depression in endangered species[14, 18]. Another question is that whether a small population is undergoing the genetic purging? Or, whether the purging in a population is effective enough to remove deleterious alleles to increase the fitness potential? Although current studies showed that the purging in some model organisms could be very effective to reduce inbreeding depression[19, 20], purging is not always existed or strong enough to alleviate the depression[4, 17, 21]. Therefore, the purging is also a key intrinsic genetic factor that was needed to be investigated for small population to help plan the genetic rescue strategies.

The Amur tiger (*Panthera tigris altaica*) is among the most endangered big cats on the planet with great symbolic significance of conservation[22] and has been prioritized for conservation for decades. The wild population has fragmented into three subpopulations, including the main one in the Sikhote-Alin Mountain containing ∼95% (415∼490) individuals, the small one in the southwest Primorsky Krai of Russia with ∼20 individuals, and a subpopulation in northeast China connecting with the two ones in Russia with ∼60 tigers[3]. Previous investigation using microsatellite markers showed that the China subpopulation has been moderately to highly inbred, suggesting a risk to lose genetic diversity rapidly[23]. This became a big concern for its sustainability. On the other hand, China has established a captive population as *ex situ* conservation resource in northeast area[24]. Population genomic analysis showed captive Amur tigers has slight inbreeding level with no apparent inbreeding depression[25]. It has been proposed recently to apply genetic rescue to ameliorate the inbreeding of the wild tigers by introducing captive genes based on the knowledge that introducing genetic variations from large and health populations can effectively improve genetic diversity and survival of small populations[11, 26, 27] and consideration of implications from successful cases[11, 28, 29]. Thus, the Amur tiger becomes an ideal model to tactically investigate the inbreeding, outbreeding, introduction of deleterious mutation and genetic purging to support future genetic rescue plans.

Although molecular markers such as mitochondrial DNA sequence and polymorphic microsatellites have been used to estimate these genetic indices, they are obviously limited in accuracy and informativeness, particularly for evaluation of inbreeding, local adaptation, mutational load and purging, which were necessary to guide reasonable genetic rescue. In contrast, whole genomic analysis can not only substantially improve the accuracy of the above-mentioned genetic indices, but also quantify the extent of inbreeding, outbreeding, purging and accumulation of mutational loads *etc*.[10, 30, 31]. Particularly, evaluating inbreeding by measuring runs of homozygosity (ROH) is highly dependent on the contiguity of reference genome, because the ROH in small populations with high degree of inbreeding is often spanning over several millions of base pairs[10, 31, 32], which can either hardly be detected or bias the data by using fragmented genomes assembled by short reads. Additionally, hidden ROHs that will be introduced into the population by future inbreeding several generations later can be also detected if the hybrid genome could be phased into haploid genomes, which is of vital importance to predict the future inbreeding. Under the background of rapid advancement of sequencing technology, decreasing cost, advanced computational power, as well as the transition from conservation genetics to conservation genomics^41, 43, 44^, [33], high-quality and haplotype-resolved reference genomes are now more crucial and urgent than ever before for supporting conservation[34].

In order to precisely reveal the intrinsic genetic factor we mentioned above in wild Amur tigers to improve the protection and conservation for this big cat, we here present a chromosome-scale and haplotype-resolved genome for a wild Amur tiger, representing the highest quality reference genome for the Amur tiger up to date. Based on the genome, we performed comparative population genomic analysis between the wild and captive Amur tigers, providing the first insights into the genetic makeup, inbreeding impacts, local adaptation, genetic load dynamics and purging effect of the wild population, providing strong implications for future genetic rescue of this species.

## 2 Results

### 2.1 Haplotype-resolved genome for the Amur tiger

We generated a high-quality and chromosome-scale HiFi genome (hereafter PtaHapG) for the Amur tiger, representing the first haplotype-resolved reference genome of the tiger. The hybrid genome size of the Amur tiger here is 2.44Gb, representing 98.75% of the estimated genome size. The contiguity of the this genome was very high with the NG50 being of 147.84 Mb, and more than 2.43 Gb (∼99.68%) scaffolds were anchored to 19 pseudochromosomes (Fig. 1A), which is consistent with the karyotypic results of the tiger and other Felidae animals [35-37]. Besides, we identified the complete X chromosome and a 10.87 Mb Y-linked region in the tiger genome by multiple lines of evidence (Fig. 1B, C). We simultaneously resulted two groups of haplotigs for the tiger genome (here after PtaHapGH1 and PtaHapGH2) (Fig. 1A). Merqury k-mer analysis showed high completeness of the haplotype-resolved genome, and meanwhile with low level of artefactual duplication (Fig. 1D). Base-level quality evaluation with QV scores, Benchmarking Universal Single-Copy Orthologs (BUSCO) analysis, genome mapping rate of short reads and structural-level accuracy evaluation all supported high completeness and accuracy of genome assembled in this study. We also found a high collinearity among the tiger and domestic cat genome, without any fissions and fusions among these three genomes. We are confident that the reference genome for the Amur tiger here is superior to most big cat genomes ever published in contiguity, completeness, and accuracy.

**Fig. 1.**
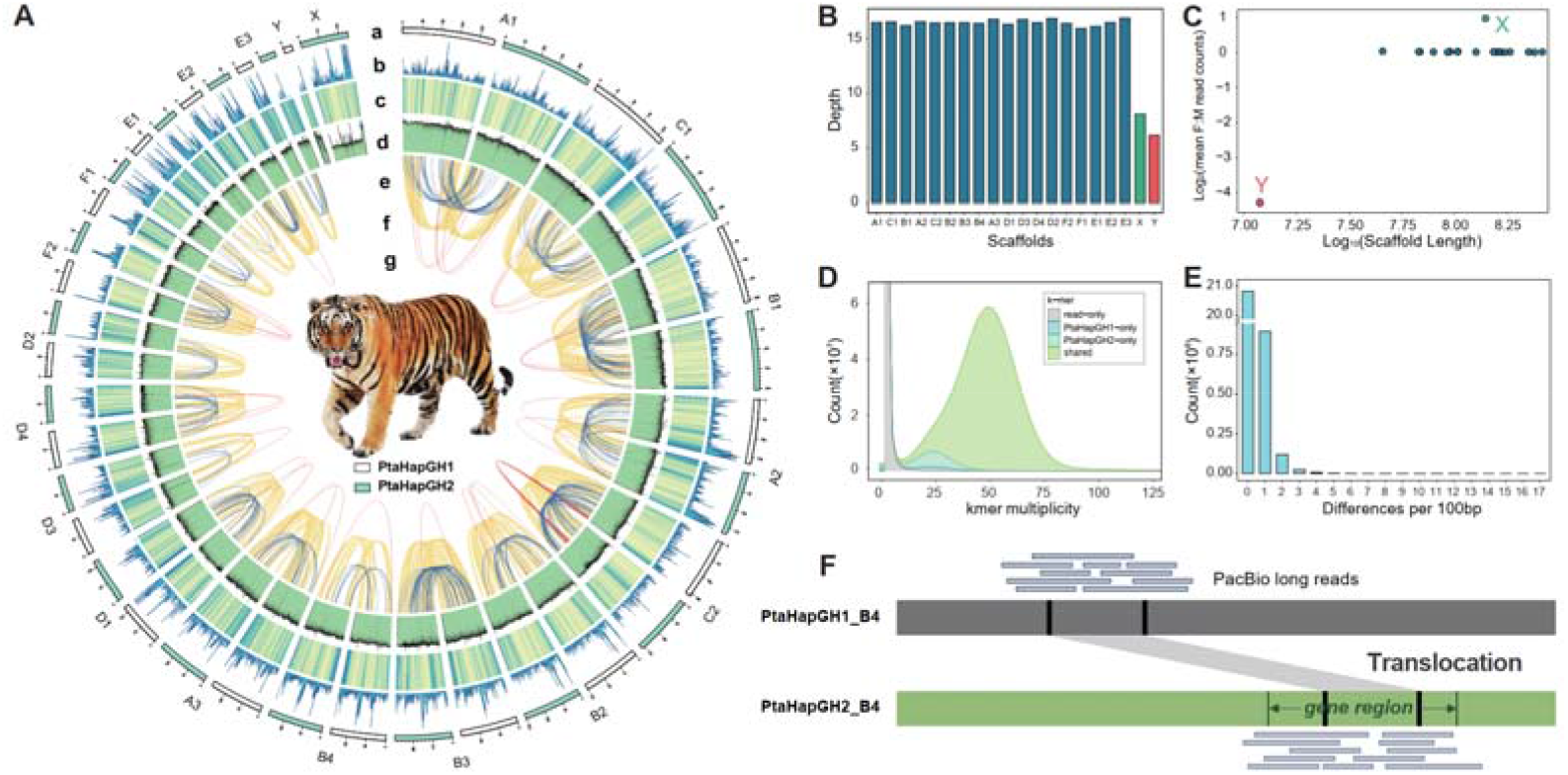
Genomic landscapes and haploid genome characteristics of the Amur tiger. A. The genomic landscape of the Amur tiger genome. a: schematic diagram of chromosomes (white: PtaHapGH1; green: PtaHapGH2); b: gene density; c: sequencing depth; d: GC contents; e: SV: duplication; f: SV: inversion; g: SV: translocation. B. Sequencing depths of the 18 autosomes, X chromosome (Hic_scaffold_10), and Y chromosome in the PtaHapG. C. Averaged depth ratios of male: female for all scaffolds in the PtaHapG. Each blue plot represents an autosome. D. K-mer spectra plot for haplotype-resolved genomes of Amur tiger produced by Merqury. Almost all haploid specific k-mers presented as single-copy (∼24X) in the genome, but the shared k-mers by the two haploid genome presented two-copy (∼49X) in the genomes. E. Pairwise differences observed in comparisons between the Amur tiger haploid genomes, the sliding window was set to be 100 bp. F. Schematic diagram of the validation of structural variates with PacBio long reads. False SV was identified if reads cannot map across the breakpoint.

We then identified 863.42 Mb (35.46%) repeat elements in the PtaHapG, with the LINE being the most abundant repeat element, making up 28.96% of the genome. We predicted 19,786 gene models in the PtaHapG by integrating *de novo* prediction, homology-based protein alignment and RNA-seq mapping evidence. Gene regions are spanning over 762.06 Mb, making up 31.29% of the genome. The BUSCO analysis showed high completeness of gene sets for all three genomes, with the lowest BUSCO score to be 95.3%. Finally, 19,771 (99.92%) genes were functionally annotated in the PtaHapG. In addition, we predicted 1128 rRNA, 1236 miRNA, 354,836 tRNA, 1647 snRNA in this genome.

### 2.2 Comparison of the two haploid genomes of the Amur tiger

In general, two haploid genomes of the PtaHapG were found to be very similar with nearly the same length, GC content, gene density, *etc*. (Fig. 1A). The single peak k-mer spectra plot and uniform reads mapping patterns to haplotype genomes indicated that we accurately generated haploid assemblies with scarce of haplotype-specific sequences (Fig. 1D). Clear one-to-one syntenic blocks between the homologous chromosome pairs between haploid genomes also showed the expected high homozygosity for the PtaHapG. The frequency of sequence differences between the two haplotypes within 100bp windows showed one peak in histogram for the PtaHapG (Fig. 1E), further indicating the high similarity between haploid genomes of the Amur tiger.

Nonetheless, we still found some chromosomal structural variants (>50bp) between PtaHapGH1 and PtaHapGH2, . In general, we found 4463 (2067 deletions, 424 duplications, 34 translocations and 1938 inversions) structural variants in the PtaHapG (Fig. 1A). All these structural variants were supported by genome mapping of both PacBio long reads and DNBSEQ short reads (Fig. 1F). 2163 genes were distributed in the structural variants of PtaHapG, with 1779 genes spanning over the breakpoints of SVs. However, we didn’t find any loss of function genes affected by SVs in the TG, further indicating the high similarity between haploid genomes.

### 2.3 Genetic differentiation between wild and captive Amur tigers

We performed whole genome resequencing of 14 wild Amur tigers collected from Jilin and Heilongjiang Province, which covers most of distribution areas of the wild Amur tiger in China (Fig. 2A). The average sequencing depth and coverage reached 31.76 folds and 97.64%, respectively. Interestingly, we found the wild and captive Amur tigers formed two distinct clades by phylogenetic tree analysis, and further supported by Admixture and PCA analysis (Fig. 2B-D). This indicated a significant genetic separation between the wild and captive Amur tigers. We further found captive individuals presented a more scattered state in the PCA result, unlike wild tigers showing a more aggregate cluster. The genetic diversity calculated from both whole chromosomal and mitochondrial genome showed a much lower evel in wild Amur tigers than in captive individuals (Fig. 2E-G).

**Fig. 2.**
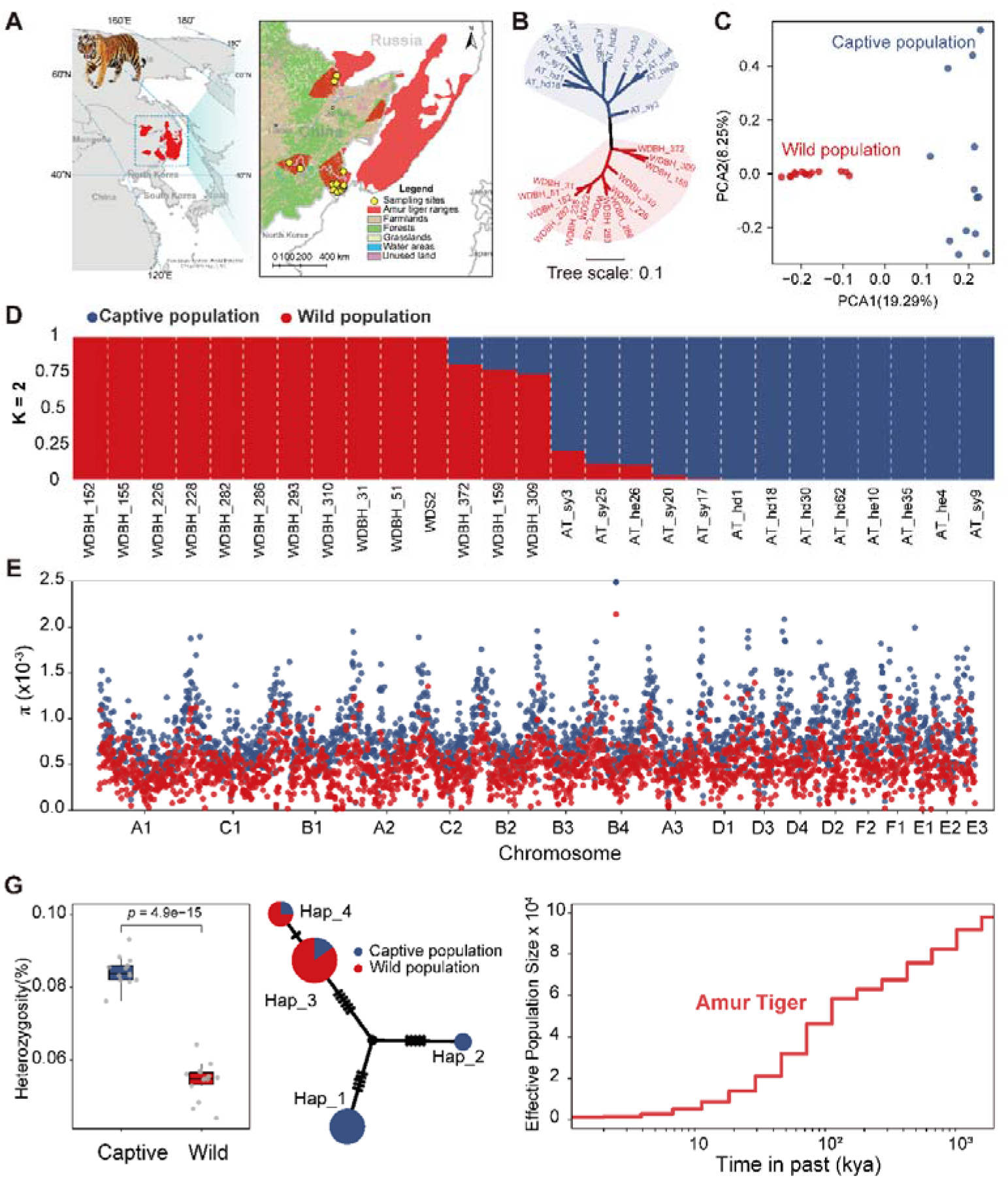
Genetic structure, genetic diversity and population demography of Amur tigers. A. The distribution area and sampling sites of wild Amur tigers in this study. B. Phylogenetic tree constructed using whole genome sequencing data of captive and wild Amur tigers. C. PCA clustering of wild and captive Amur tigers. D. Genome-wide admixture analysis of captive and wild Amur tigers. E. Genome-wide genetic diversity (π) in captive and wild Amur tigers. F. Comparison of genome-wide heterozygosity between captive and wild Amur tigers. G. Network analysis of mitochondrial haplotypes of Amur tigers. H. The dynamic of effective population size in the evolutionary history of wild Amur tiger.

### 2.4 Recent demographic history of the wild Amur tiger

To investigate the historical dynamics of population demography of wild Amur tigers, we inferred and described the changes of effective population size (*N*_*e*_) over time. In general, *N*_*e*_ for the Amur tiger has been declining over its entire evolutionary history (Fig. 2H). However, we did not observe any increase of *N*_*e*_ for wild Amur tiger throughout the recent 10,000 years as for the slight rebound in captive tigers[25]. In addition to low genetic diversity (Fig. 2E, F), the continuous population declining once again reminds us the latent extinction risk of wild Amur tigers.

### 2.5 Estimates of inbreeding in wild Amur tigers

Here we quantified inbreeding in wild Amur tigers by screening genome-wide ROHs, which is a key genetic indicator to guide possible genetic rescue[25]. We found a total of 26,198 ROHs longer than 100kb, averaged 254.19 Kb in length, and the longest ROH reached 35.11 Mb. The averaged number and length per wild individual were 1871 ± 84 Mb and 1160.73□±24.80 Mb, respectively. For long ROHs originated from recent inbreeding, we found 3746 ROHs longer than 1Mb in the wild population with averages per-individual of 267 ± 11 in number and 696.92 ± 44.56Mb in length. Similarly, a total of 368 ROHs longer than 5Mb were identified from this population with averages of 26 ± 4 in number and 213.61 ± 37.10Mb in length.

We further measured the inbreeding level in wild Amur tigers using F_ROH_ by comparing with captive individuals. In general, the level of F_ROH_ was found to be negatively related with genome-wide heterozygosity in both captive and wild Amur tigers, but the inbreeding level in the wild Amur tiger is higher than captive individuals (Fig. 3A). The F_ROH>100kb_, F_ROH>1Mb_, and F_ROH>5Mb_ were 0.51±0.01, 0.30±0.02, and 0.09±0.02 per individual, respectively, in the wild population, while in captive Amur tigers, we showed 0.26±0.01, 0.17±0.01, and 0.08±0.01 for the same order of F_ROH_ (Fig. 3B). It is evident that the wild population is more inbred than captive tigers by almost two times as measured with F_ROH>100kb_. The fraction of inbreeding introduced by the recent ancestor (up to 5 generations ago)[4] was fairly milder in the wild population, but the F_ROH>5Mb_ was still 1.13 times higher than in captive tigers, although this difference is not significant (Fig. 3C). In addition, 95% and 64% ROH fragments were distributed in the IBD regions in the wild and captive Amur tigers, indicating that most of the ROHs were originated from the inbreeding, especially in the wild population. Overall, the inbreeding level in wild individuals were significantly higher than that in captive Amur tigers (*p*=5.6e-15) (Fig. 3B, D). This was also supported by less mitochondrial haplotype diversity in the wild population (Fig. 2G).

**Fig. 3.**
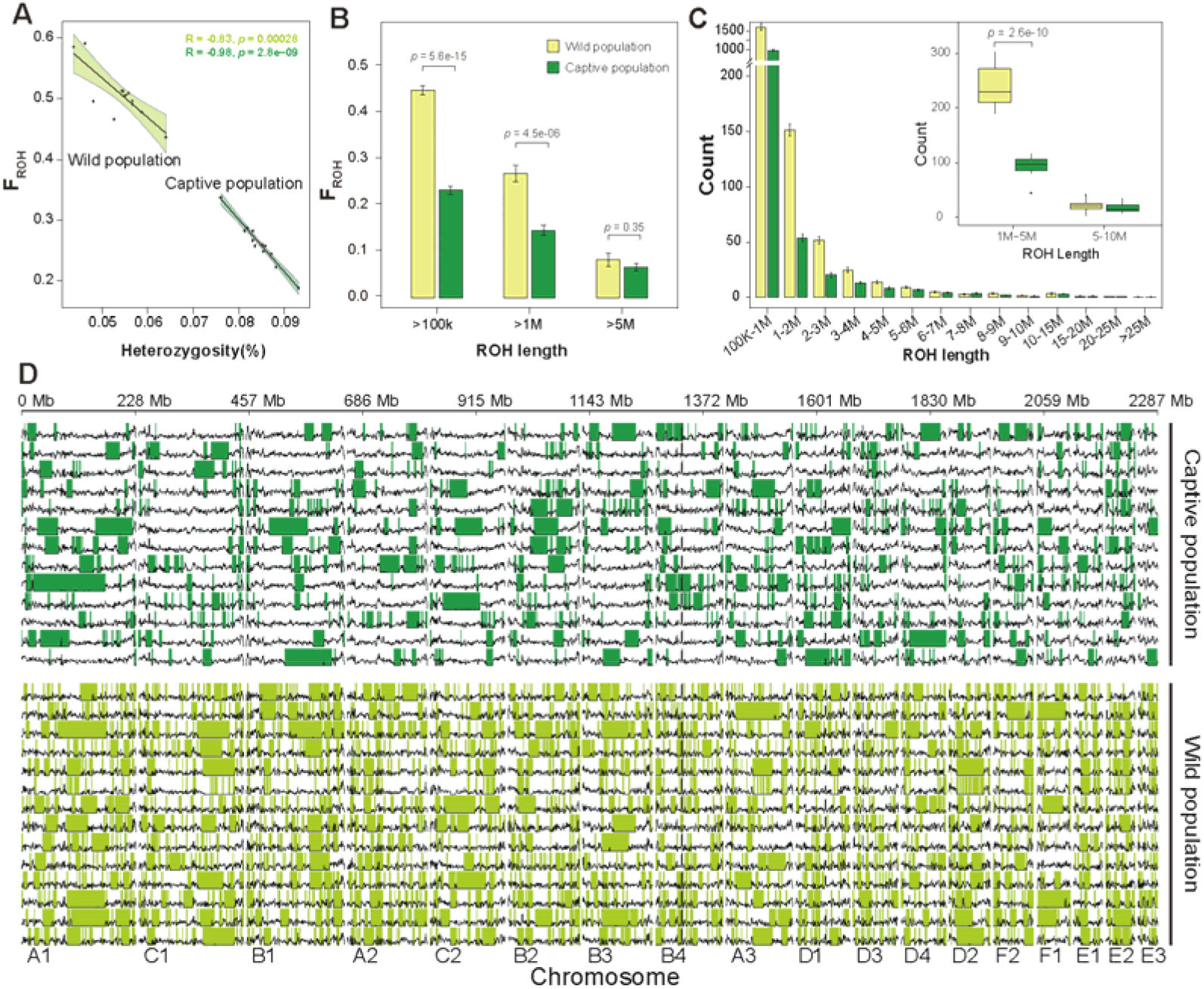
Genome-wide inbreeding estimation of Amur tigers. A. A positive relationship between the heterozygosity and F_ROH_ in Amur tiger. The wild Amur tigers was represented in light green and the captive tigers was in green. B. The comparison of averaged F_ROH_ in wild and captive Amur tigers. C. The length distribution of ROH fragments in Amur tigers. The wild Amur tiger showed a higher level ROH less than 5Mb. D. A fine-scale distribution of ROH larger than 1Mb in wild and captive Amur tigers.

To look into detailed inbreeding history of wild Amur tigers, we dissected the ROH distribution by length. We found the large proportion of ROH fragments were restricted to < 1Mb (Fig. 3C), indicating that inbreeding has occurred since 26 generations ago (based on a recombination rate of 1.9cM/Mb from the domestic cat[39]). We further observed that the number of ROHs shorter than 5Mb was much greater in the wild Amur tigers (1845 ± 87.11) than in captive tigers (1071.46 ± 31.55, *p*=2.7e-07), signifying that the wild population has experienced progressive inbreeding five generations ago. However, the inbreeding levels in the recent five generations (ROH > 5M) was not significant (*p*=0.26) between the wild (26.29 ± 3.81) and captive tigers (20.69 ± 2.61) although the wild population presented more ROHs. To reveal the inbreeding occurred as recent as the past three generations, we continued focusing on ROH larger than 10Mb. Results showed that the average F_ROH>10Mb_ was 3% ± 1% in the wild Amur tigers, much lower than that in the South China tiger (*P. t. amoyensis*) (6%)[25] and the small isolated Bengal tiger (*P. t. tigris*) population in India (28%)[4]. This suggests the negative effects signified by recent inbreeding may not be that serious in the wild Amur tigers so far.

### 2.6 Detection of genome-wide mutational loads in wild Amur tigers

Mutational load is a kind of intrinsic threat to the survival and fitness of endangered species. Here, we firstly investigated the genome-wide mutational loads in protein-coding regions in wild Amur tigers by focusing on deleterious non-synonymous SNPs (dnsSNPs) and loss of function mutations (LOFs), which are both expected to potentially affect the gene functions. Totally, we found 562 and 130 dnsSNPs (Grantham Scores ≥150) and LOFs in the wild population, which is lower than that in captive Amur tigers, particularly for the LOF with a much greater difference (Fig. 4A, 5A). We obtained the same result when only considered the derived LOFs and dnsSNPs. Similar result was observed when we used Genomic Evolutionary Rate Profiling (GERP) method to estimate the relative mutational load by measuring the number of derived alleles in the strict evolutionary constraints regions. We further compared the wild and captive population by using the Rxy method to estimate whether the excess of deleterious derived alleles existed in wild Amur tigers. The results showed reduced LOF and missense mutations, but increased synonymous mutations in the wild population (Fig. 4B). In addition, the frequency of LOFs and dnsSNPs inside ROH regions are much lower than that in non-ROH regions in both wild and captive Amur tigers, indicating the purging has occurred in both populations (Fig. 4C, D). Moreover, the frequency ratios of LOFs and dnsSNPs inside ROH to those outside ROH were both much lower in the wild population than in captive Amur tigers (Fig. 4E). This suggests that higher inbreeding in the wild population facilitated purging of deleterious mutations with greater stringency. In addition, we observed a higher proportion of derived LOFs were in homozygous state in the wild populations than that in captive Amur tigers, but we didn’t observe significant differences between two populations for the dnsSNPs (Fig. 4F). High proportion homozygous LOFs indicated that fitness cost result from LOFs is higher in the wild population, despite of the existence of purging.

**Fig. 4.**
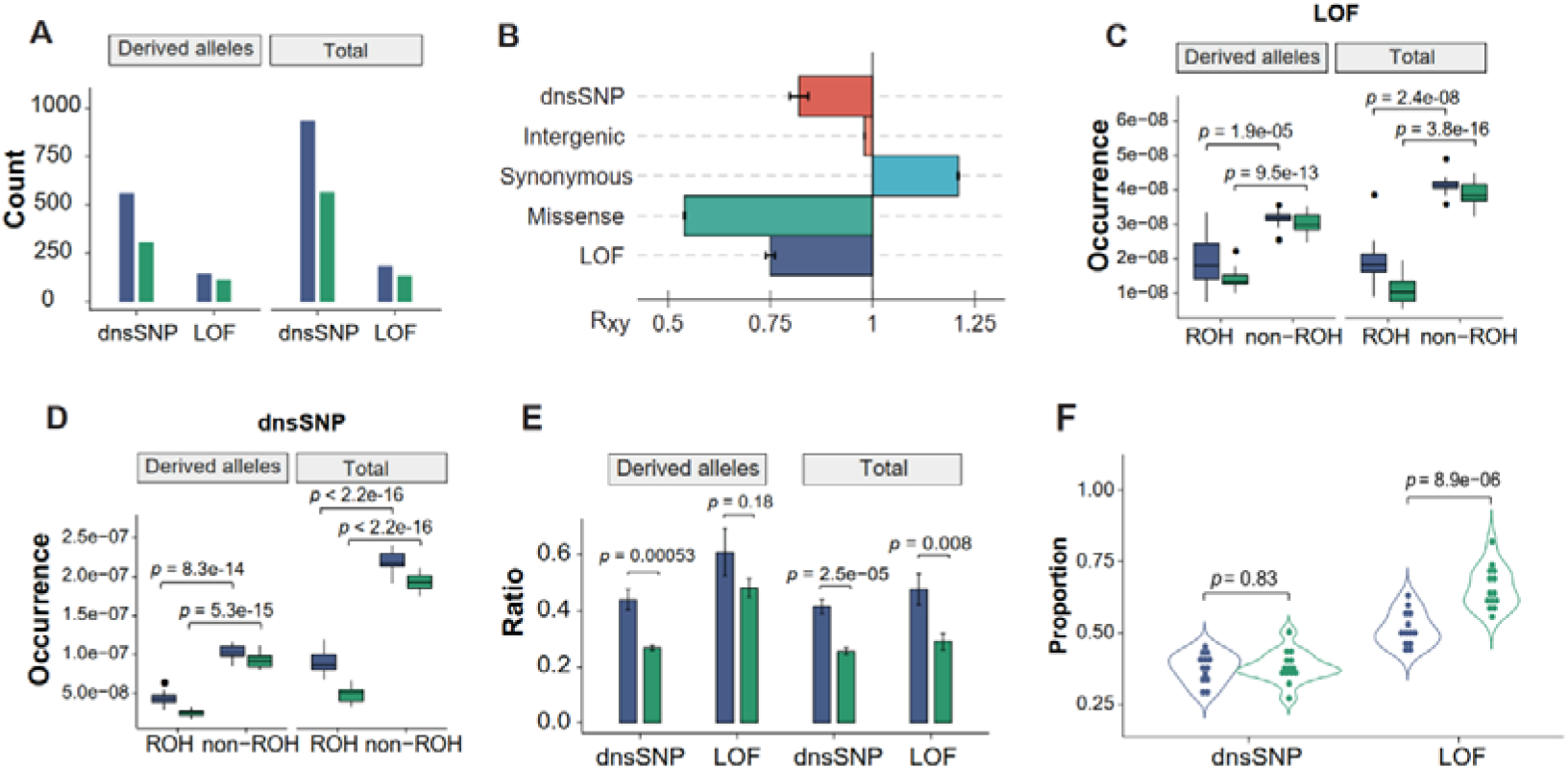
Characteristics of mutational load in captive and wild Amur tigers. A. Comparisons of the number of mutational loads for both LOF and dnsSNP in captive and wild Amur tigers. B. The Rxy ratio between derive alleles in wild (x) and captive (y) for dnsSNP, intergenic regions, synonymous, missense and LOF. The Rxy <1 indicated the population y has more derived alleles than population x. C. The comparison of frequency for LOF variants (LOF number / Total base number of ROH or non-ROH genomic regions) between inside and outside of ROH regions. D. The comparison of frequency for dnsSNP variants between inside and outside of ROH regions. E. The ratio of frequency for LOF/dnsSNP inside ROH to that outside ROH in wild and captive Amur tigers. This ratio in the wild Amur tigers was significantly higher than captive Amur tigers. F. The proportion of homozygous LOF and dnsSNP in wild and captive Amur tigers. For A, C-F, the blue represented the captive Amur tiger and the green presented the wild Amur tiger.

To further dissect the effects of nature selection and random genetic drift on purging of deleterious mutations in the tiger populations (Fig. 5A), we performed the Site Frequency Spectra (SFS) analysis. We found the number of derived neutral alleles fixed in wild (0.15) and captive Amur tigers (0.05) were all much greater than damaging alleles (wild: 0.0014, captive: 0.0027), while the frequency of fixed alleles was much higher in the wild population (∼0.15) (Fig. 5B). For polymorphic loci, the SFS was flat in the wild population but exhibited a continuing decline in captive Amur tigers, suggesting captive Amur tigers harbored more rare derived alleles. These above-mentioned genetic features indicated a strong bottleneck in the wild population compared with the captive individuals. Furthermore, it could be expected that the deleterious alleles in the population should be at low frequency given natural selection were more effective to eliminate deleterious alleles than neutral ones. In this study, we found that, in the wild population, the average frequency of deleterious alleles was 0.14±0.002, significantly lower than that of neutral alleles (0.39±0.00021, *p*=2.2e-16), especially for the deleterious allele with a frequency larger than 0.3. The frequency spectra were much more flat and lower in the wild population. From results above we inferred that the purifying selection contributed to the purging of mutational loads in the wild Amur tigers.

**Fig. 5.**
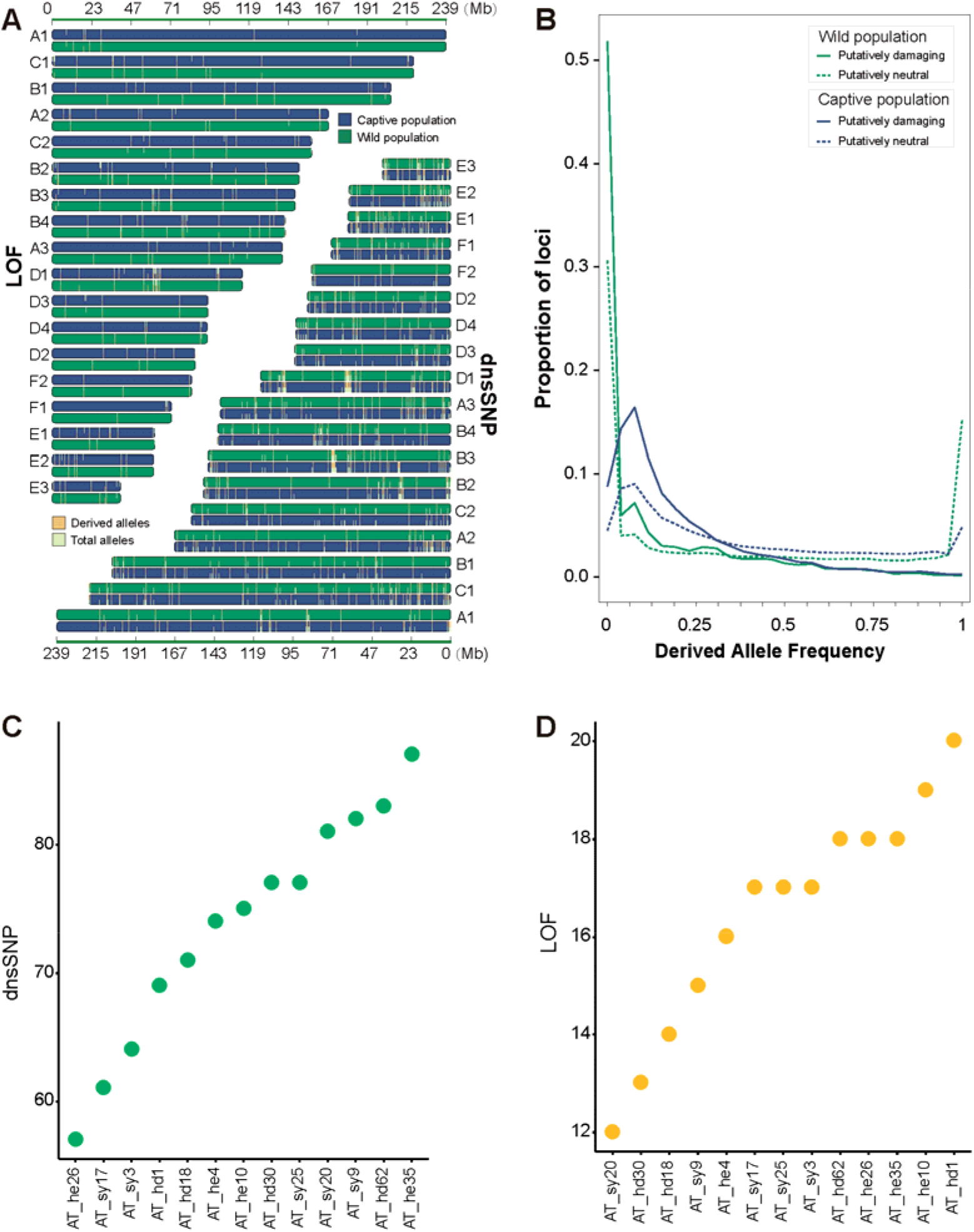
Results of dissecting natural selection on the purging of deleterious mutations and the distribution of deleterious mutations across the whole genome. A. Distribution of LOF and dnsSNP across each chromosome in wild and captive Amur tiger population. B. Site-frequency spectrum for damaging and neutral mutations for each population. C. Potentially new dnsSNP mutations introduced to the wild population by captive individuals if gene flow occurred. D. Potentially new LOF mutations introduced to the wild population by captive individuals if gene flow occurred.

We finally predicted the possibly introduced deleterious mutations from the captive to the wild population by simulating pairing regimes between the two populations. We found 315 and 67 derived dnsSNPs and LOFs would be introduced into the wild population when genetic rescue is applied by using the current captive population. However, the introduced captive deleterious mutations vary depending on individual gene donors (Fig. 5C, D).

### 2.7 Genomic signatures of local adaptation in Amur tigers

Local adaptation is a key factor should be taken into consideration when making conservation strategies for endangered species[15, 40, 41]. Here we applied Cross Population Extended Haplotype Homozygosity (XP-EHH) method to screen the possible signals of recent positive selection in the captive Amur tigers compared to the wild population to facilitate the genetic rescue by reintroduction of captive individuals to the wild. We found 4559 and 4516 SNPs were identified under strong positive selection in the wild and captive Amur tigers, respectively, identified by setting a XP-EHH score in the top 0.1%. Only 56 and 63 genes were found to be under recently positive selection in the wild and captive Amur tigers, respectively. In the wild population, we found many GO terms were closely related with immunity, defense response, and response to environmental pollutants. We also found a KEGG pathway, Toll-like receptor signaling pathway, was significantly enriched in the wild population. However, no evidence was found for overrepresented GO terms and KEGG pathways in captive Amur tigers.

## 3. Discussions

### 3.1 Improved reference genome for the Amur tiger

Currently, the combination of high-fidelity (HiFi) reads and HiFi-specific assembler can generate high-quality and haplotype-resolved *de novo* assembly, representing one of the most promising strategy for genome assembling by far[42, 43]. The reference genome we report here is the first haplotype-resolved reference genome for the tiger[44]. There has been reported two Amur tiger genome assemblies, including a second-generation genome (PanTig1.0) [45, 46] and a chromosome-scale genome (PanTig2.0) assembled by PacBio long reads. The Amur tiger genome assembled in this study have a much higher contiguity, with a contig N50 of 26.97 Mb, which is ∼692-fold and ∼3-fold longer than the PanTig1.0 (N50: 0.039 Mb) and PanTig2.0 (N50: 9.52 Mb), and the contig number of PtaHapG (259) much fewer than PanTig2.0 (3117). This significant improvement in contiguity will greatly improve the evaluation of inbreeding in tigers. In particular, by comparing this genome to our previously published chromosome-scale genome of a captive Amur tiger, we found 37.73 Mb wild individual specific sequences harboring the gene OR56A3 which was missing in the captive genome, further indicating the superiority of the HiFi genome we assembled here.

### 3.2 Mutational load purging in wild inbred Amur tigers

The wild population of Amur tiger has fragmented into three subpopulations (Fig. 2A). The subpopulation in northeast China connecting with the two ones in Russia restored ∼60 tigers at present from ∼26 in 2014[3]^,[47]^ . Previous study showed a risk of this population to lose genetic diversity rapidly due to moderate to high inbreeding[23]. Thus, inbreeding became a big concern for its sustainability. On the other hand, China has established a captive population as *ex situ* conservation resource in northeast area[24]. Population genomic analysis showed moderate inbreeding level for captive Amur tigers. To draw a general picture of genetic status of the wild population in China, we used advantage of our haplotype-resolved and super high contiguity resolution genome PtaHapG to precisely estimate and predict inbreeding level and genomic consequence for the representatives of the captive and wild Amur tigers (Fig. 3D).

We show the inbreeding levels (F_ROH_) inferred from ROHs longer than 100 kb, 1Mb and 5Mb were all higher in wild tigers than in captive tigers by 1.96, 1.76 and 1.13 times (Fig. 3B). Overwhelming proportion (∼95%) of ROHs (>100kb) distributed in IBD regions in the wild population, suggesting majority of ROHs were resulted from inbreeding, although shared bottleneck history may still contribute to the genome homozygosity[48]. Furthermore, majority of ROH fragments were restricted to < 1Mb, suggesting inbreeding could be dated back to 26 generations ago (estimated based on a recombination rate of 1.9cM/Mb from the domestic cat[39]). A relative larger number of ROHs < 5Mb were observed in wild tiger genomes than in captive genomes, signifying the wild population has experienced more inbreeding five generations ago. The average F_ROH_>10Mb was half the South China tiger (*P. t. amoyensis*) (6%) and one tenth the small isolated Bengal tiger (*P. t. tigris*) population in India (28%)[4], suggesting the negative effects signified by recent three generations may not be that serious so far. However, the number of ROHs shared among tigers in the wild population was much higher than that in captive Amur tigers, further suggesting more intensive inbreeding in wild Amur tigers. Inbreeding for such fragmented population[49] will doubtlessly lead to loss of genetic diversity driving it towards extinction without timely human interference.

Fortunately, we found the mutational loads in wild Amur tigers, including both LOF and dnsSNP variants, are much lower than that in the captive tigers (Fig. 4A). Expectedly, this might be resulted from purging promoted and maintained by random genetic drift and/or the combination of inbreeding and purifying selection. The wild Amur tigers has experienced sharp decline within the most recent 1000 years leaving a small *Ne* (Fig. 2H). Recent decades of isolation would further reduce the *Ne*. Genetic drift in this case might have effectively acted on this population leading to random loss or fixation of deleterious alleles with equal probability [10]. However, we detected an elevated frequency of fixed neutral alleles comparing to damaging alleles in the wild population, suggesting purifying selection may have been acting on the wild Amur tiger population to reduce the inbreeding depression by purging of damaging alleles. Furthermore, the LOF mutations were much less than dnsSNP (Fig. 4A), and the occurrence of LOF and dnsSNP was significantly more frequent outside ROH regions than that inside ROH regions (Fig. 4C, D). This suggests that homozygous deleterious mutations were more likely purged than heterozygous ones, and purging was stronger for LOFs that are more recessive than missense mutations.

In general, our genomic analysis showed the inbreeding of the wild Amur tiger population in northeast China could be dated back to at least 26 generations ago. Fortunately, up to the most recent generations inbreeding has not yet reached the level to express severe inbreeding depression. Inbreeding combining with purifying selection has facilitated mutational load purging, but meanwhile presents a risk to lose genetic diversity. In contrast, captive Amur tigers had also been inbred 26 generations ago but experienced relative lower inbreeding in the past five generations. Mutational load purging in the captive Amur tigers is not so effective as in the wild population, very likely due to effective husbandry management[51].

### 3.3 Implications for genetic rescue of the wild population

The above analysis shows inbreeding had taken place and developed to moderate to high level in the wild Amur tiger. Although the population number of wild Amur tiger has restored successfully in the past decade (National Forestry and Grassland Administration, P. R. China, 2022), the security is still worrying and ameliorating inbreeding should be taken into the main goals of next phase conservation.

One immediate and necessary approach is to build ecological corridors to connect up population patches within China and with Russian populations to eliminate landscape resistance of migration and improve genetic communications[3]. This is key infrastructure supporting long-term survival[52]. The second and parallel approach is genetic rescue using the healthier captive population[53]. We found that captive tigers are genetically distant from the wild ones and higher in genetic diversity (Fig. 2B-G), which are very potential to ameliorate ROHs in the wild genomes. However, we also found that captive Amur tigers carries greater number of mutational loads (Fig. 4A) and a considerable proportion are absent in the wild population (Fig. 5B). This suggests introducing captive genes into the wild population is risky to simultaneously introduce novel mutational loads that may make further negative impacts. Therefore, cautions should be made to avoid this side-effect.

In this study, we established partial list of mutational loads for the wild and captive tigers and predicted 315 derived dnsSNPs and 67 LOFs likely to be introduced into the wild population by simulating currently studied captive tigers for genetic rescue (Fig. 5C, D). The introduced mutations vary depending on gene donors, leaving possibilities for selecting the optimal candidate tigers. Our study provides an instance to accurately evaluate the extent to which the ROHs on wild genomes could be ameliorated, mutational loads be introduced, and the likelihood the introduced loads could be subjected to remission by heterozygote under possible reproductive regimes. For planning real genetic rescue, the list of mutational loads should be completed by sampling all, at least majority of wild tigers and all potential captive gene donors. The haplotype resolution reference genome PtaHapG provides a firm platform to carry out elaborate inference in this regard.

## Acknowledgments

This study was supported by the Fundamental Research Funds for the Central Universities of China (2572022DQ03), National Natural Science Foundation of China (32170517) and the Guangdong Provincial Key Laboratory of Genome Read and Write (grant No. 2017B030301011). This work was also supported by China National GeneBank (CNGB).

## Author Contributions

These authors contributed equally: Tianming Lan, Haimeng Li, Le Zhang, Minhui Shi, Boyang Liu. Yanchun Xu, Huan Liu, Guangshun Jiang and Tianming Lan initiated and designed the project. Dan Liu, Yue Zhao, Weiyao Kong, Yue Ma, Boyang Liu and Le Zhang collected the samples. Qing Wang, Jiangang Wang, Xinyu Wang, Yuxin Wu, Jiale Fan, Xiaotong Niu and Liangyu Cui performed the RNA/DNA isolation. Haorong Lu, Shaofang Zhang, Jieyao Yu performed the DNA libraries construction and sequencing. Nicolas Dussex and Tianming Lan coordinated the data analysis. Haimeng Li, Minhui Shi, Le Zhang and Boyang Liu carried out the data analysis. Tianming Lan, Haimeng Li, Minhui Shi, Boyang Liu and Yanchun Xu wrote the manuscript. Guangshun Jiang, Shanlin Liu, Love Dalén, Nicolas Dussex and Yanchun Xu revised the manuscript. Guangshun Jiang, Huan Liu, and Yanchun Xu provided the supervision of this project.

## Data and Code Availability

The data that support the findings in this study have been deposited into CNGB Sequence Archive (CNSA) [54]of China National GeneBank DataBase (CNGBdb) [55] with accession number CNP0003803.

